# Silencing KCC2 in mouse dorsal hippocampus compromises spatial and contextual memory

**DOI:** 10.1101/2022.02.18.481031

**Authors:** Clémence Simonnet, Manisha Sinha, Marie Goutierre, Imane Moutkine, Stéphanie Daumas, Jean Christophe Poncer

## Abstract

Delayed upregulation of the neuronal chloride extruder KCC2 underlies the progressive shift in GABA signaling polarity during development. Conversely, KCC2 downregulation is observed in a variety of neurological and psychiatric disorders often associated with cognitive impairment. Reduced KCC2 expression and function in mature networks may disrupt GABA signaling and promote anomalous network activities underlying these disorders. However, the causal link between KCC2 downregulation, altered brain rhythmogenesis and cognitive function remains elusive. Here, by combining behavioral exploration with *in vivo* electrophysiology we assessed the impact of chronic KCC2 silencing in mouse dorsal hippocampus and showed it compromises both spatial and contextual memory. This was associated with altered hippocampal rhythmogenesis and neuronal hyperexcitability, with increased CA1 pyramidal cell burst firing during non-REM sleep. Reducing neuronal excitability with terbinafine, a specific Task-3 leak potassium channel activator, occluded the impairment of contextual memory upon KCC2 silencing. Our results establish a causal relationship between KCC2 expression and cognitive performance and suggest that impaired rhythmopathies and neuronal hyperexcitability are central to the deficits caused by KCC2 silencing in the adult mouse brain.

## Introduction

Fast inhibitory synaptic transmission in the central nervous system relies on chloride conductance associated with both GABAA and glycine receptors. Whereas most cell types display high intracellular chloride concentration ([Cl]_i_), mature neurons maintain much lower [Cl]_i_ and thereby ensure a hyperpolarizing chloride influx upon GABAA or glycine receptor activation (Kaila *et al*, 2014a). This property is supported by the expression of a neuron-specific chloride extruder, the potassium-chloride co-transporter KCC2. Early expression of the sodium-potassium-chloride cotransporter NKCC1, a ubiquitous chloride importer, and delayed upregulation of KCC2 expression combine to produce a developmental shift in neuronal [Cl]_i_ and, consequently, in the polarity of GABA signaling in the developing CNS, from excitatory to inhibitory or shunting (Kaila *et al*., 2014a; Rivera *et al*, 1999; Virtanen *et al*, 2021).

However, in the mature brain, KCC2 may be downregulated in a variety of pathological conditions, including brain trauma (Bonislawski *et al*, 2007; Sawant-Pokam *et al*, 2020), stroke (Jaenisch *et al*, 2010), stress (MacKenzie & Maguire, 2015; Sarkar *et al*, 2011), epilepsy (Huberfeld *et al*, 2007; Pallud *et al*, 2014; Pathak *et al*, 2007; Rivera *et al*, 2004) as well as schizophrenia (Arion & Lewis, 2011; Hyde *et al*, 2011; Sullivan *et al*, 2015) and autism spectrum disorders (Banerjee *et al*, 2016; Bertoni *et al*, 2021; Duarte *et al*, 2013; Tang *et al*, 2019). Most of these disorders are associated with cognitive impairment, including episodic and working memory deficits (Dillon & Pizzagalli, 2018; Dupont *et al*, 2000; Gur & Gur, 2013; Lim & Alexander, 2009). However, whether and how KCC2 downregulation in pathology may contribute to memory deficits has remained unexplored.

Downregulated KCC2 expression is classically thought to cause a pathological shift in the polarity of GABA signaling, from inhibitory to excitatory. Thus, depolarizing GABAA receptor-mediated responses are observed in most aforementioned conditions. Such excitatory actions of GABA in mature neuronal networks may then promote anomalous network activities underlying pathological symptoms (Banerjee *et al*., 2016; Buchin *et al*, 2016; Kaila *et al*, 2014b; Pallud *et al*., 2014; Tyzio *et al*, 2014). However, KCC2 downregulation may also affect neuronal function independent of GABA signaling. KCC2 was shown to regulate activity-dependent synaptic delivery of AMPA receptors through interaction with actin-related protein partners (Chevy *et al*, 2015; Gauvain *et al*, 2011; Li *et al*, 2007; Llano *et al*, 2015), thereby gating long term potentiation induction (Chevy *et al*., 2015; Chevy *et al*, 2020; Virtanen *et al*., 2021). In addition, KCC2 was shown to control membrane trafficking of leak potassium channels Task-3 through molecular interaction. KCC2 knockdown therefore induces neuronal and network hyperexcitability by reducing Task-3 channel membrane expression and function (Goutierre *et al*, 2019). These effects may combine to perturb both long term synaptic plasticity and cortical rhythmogenesis, two processes critically involved in hippocampus-dependent memory encoding and consolidation (Buzsaki, 2015; Colgin, 2016; Nicoll, 2017; Whitlock *et al*, 2006).

Here we tested whether KCC2 downregulation in mouse dorsal hippocampus was sufficient to impair memory and explored the underlying mechanisms. Using RNA interference to knockdown but not fully ablate KCC2 expression in a cell-type specific manner, we observed deficits in both spatial and contextual memory upon KCC2 silencing in hippocampal principal neurons but not GABAergic interneurons. Our data indicate this effect is associated with altered hippocampal rhythmogenesis and involves, at least in part, neuronal hyperexcitability and bursting behavior.

## Results

### KCC2 down-regulation in dorsal hippocampal neurons impairs spatial and contextual memory in mice

Genetic ablation of *Slc12a5* encoding the KCC2 transporter in mice is associated with severe developmental defects leading to premature death around birth, due to respiratory complications (Hübner et al., 2001) and seizures (Uvarov et al., 2007). In order to evaluate the impact of KCC2 downregulation, as observed in the human pathology, independent of developmental defects associated with constitutive ablation, we used virus-based chronic silencing by RNA interference. Adult mice were infected bilaterally in dorsal hippocampus with AAV vectors expressing either non-target (shNT) or previously validated KCC2-specific (shKCC2) shRNAs (Bortone & Polleux, 2009) (Figure 1A-C). Effective KCC2 silencing was verified after 2-4 weeks by immunostaining (Figure 1D) and western blot analysis of micro-dissected hippocampal extracts (Figure 1E-F). KCC2 expression was reduced by >70% in hippocampal extracts from mice infected with virus expressing shKCC2 as compared to shNT (t-test p<0.001). Viral infection *per se* yielded no change in KCC2 expression, as no significant difference was detected between samples from mice infected with control (shNT) virus and non-infected animals (t-test p=0.388).

**Figure 1.**
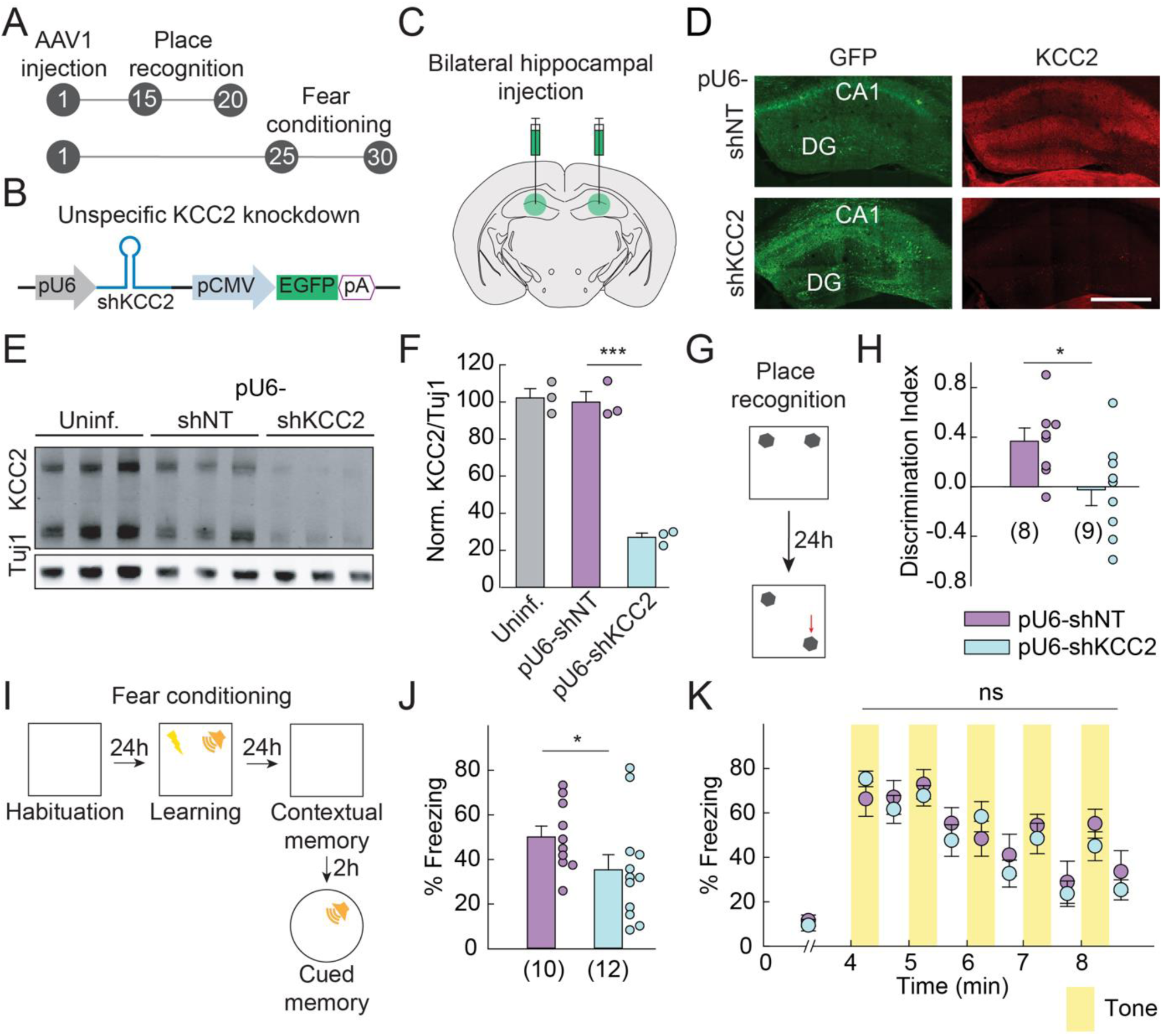
KCC2 down-regulation in dorsal hippocampus impairs spatial and contextual memory. **A.** Timeline of the experiment (days). **B.** Design of the viral vector for unspecific KCC2 knockdown. **C.** Wild-type mice were injected bilaterally in the dorsal hippocampus with AAV vector designed to knockdown KCC2. **D.** Representative confocal maximal projection images of hippocampal coronal sections immunostained for GFP and KCC2 from mice infected with U6-shNT or U6-shKCC2-expressing AAV vectors, showing massive KCC2 knockdown throughout hippocampal fields in the later. Scale: 500 µm. **E.** Representative immunoblot of KCC2 in hippocampal protein extracts from non-infected mice (n=3) and mice infected either with U6-shNT (n=3) or U6-shKCC2 (n=3) expressing viruses. **F.** Quantification of KCC2 expression, relative to that from mice infected with U6-shNT vector (t-test, p<0.001). Tubulin was used as an internal standard. **G.** Protocol of the place recognition task. **H.** Discrimination index showing control mice learned to identify the moved object better than KCC2-knockdown mice (t-test, p=0.017). **I.** Protocol of the fear conditioning task. **J.** In the contextual memory test, freezing was assessed during the first 3 minutes of exploration of the foot-shock associated cage. Mice infected with U6-shKCC2 vector display reduced freezing time compared to mice infected with U6-shNT vector (t-test, p=0.050). **K.** Summary graph showing the time course of freezing upon exposure to foot-shock associated sound. Cued memory retention shows no difference between both groups.

We next explored the effects of chronic KCC2 downregulation on mouse behavior. No consistent difference was observed between control and KCC2 knockdown mice in terms of locomotor activity or anxiety (Figure EV1). Thus, we tested whether KCC2 silencing might affect hippocampus-dependent spatial (Barker & Warburton, 2011) and contextual memory. In an object-place recognition paradigm (Figure 1G), mice explored two identical objects for 10 minutes in an arena with a cue card, and one of the objects was moved to a new location in the arena 24 hours later. While control mice showed a significant preference for the moved object (Figure 1H), this preference was significantly reduced in KCC2-knockdown mice (t-test p=0.017). This shows that KCC2 silencing in dorsal hippocampus compromises spatial memory. We next evaluated the performance of mice in contextual and cued fear memory. Mice received 4 foot-shocks associated with a tone on the training day. On the next day, contextual and cued memories were tested (Figure 1I). In the hippocampus dependent contextual memory test, the freezing response of KCC2-knockdown (shKCC2) mice was significantly reduced compared to that of control (shNT) mice (Figure 1J; t-test p=0.050). However, no difference in freezing behavior was observed between the two groups in the sound-cued memory test (Figure 1K ; two-way repeated measures ANOVA p=0.329), which primarily depends on the amygdala and not the hippocampus (Phillips & LeDoux, 1992). Our results therefore suggest that chronic KCC2 silencing in the dorsal hippocampus specifically impairs both spatial and contextual memory.

### KCC2 silencing in hippocampal principal neurons is sufficient to compromise contextual memory

We next aimed to explore the cellular and network mechanisms underlying memory deficits upon KCC2 knockdown in dorsal hippocampus. Since KCC2 is expressed in both hippocampal principal cells and at least some interneurons subtypes (Gulyas *et al*, 2001; Otsu *et al*, 2020), we tested whether KCC2 silencing in either neuron subtype might be sufficient to recapitulate memory deficits induced by unspecific silencing. In order to specifically suppress KCC2 expression in principal neurons or interneurons, shRNA sequences were imbedded into a mir30 backbone (Fellmann *et al*, 2013) and inserted downstream Pol II promoters specific to either cell type in the forebrain (pCamKII (White *et al*, 2011) and pmDlx (Dimidschstein *et al*, 2016), respectively )(Figure 2B). These constructs efficiently suppressed KCC2 expression in principal neurons and interneurons of dorsal hippocampus, respectively, as illustrated by immunofluorescence detection (Figures 2C-D). Five to six weeks following viral injection, mice were then exposed to a fear-conditioning paradigm (Figure 2A). During the contextual memory test, mice infected with a virus expressing pCaMKII-shmirKCC2 targeting principal neurons showed significantly less freezing than control mice (t-test p=0.019; Figure 2F), comparable to the effect observed in mice with cell-type unspecific silencing (Figure 1J). In contrast, mice infected with a virus expressing pmDlx-shmirKCC2 targeting GABAergic interneurons displayed a freezing response comparable to that of the control group (t-test p=0.215). Again, mice expressing either pCaMKII-shmirKCC2 or pmDlx-shmirKCC2 in the dorsal hippocampus showed no deficit in cued memory as compared to their respective controls (Figure 2G).

**Figure 2.**
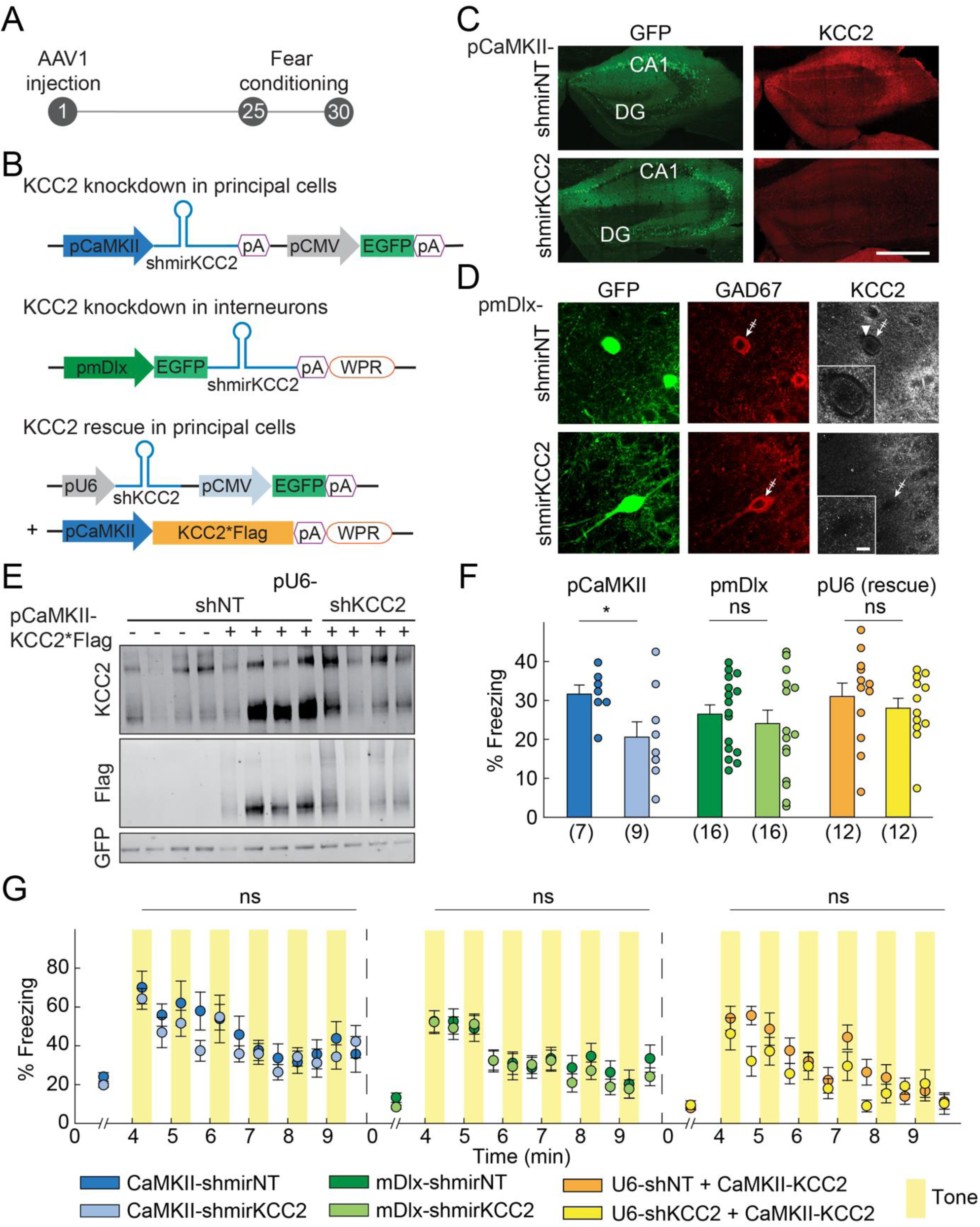
KCC2 down-regulation in principal cells is sufficient to alter contextual memory. **A.** Timeline of the experiment (days). **B.** Viral vectors used to knockdown KCC2 in principal cells or interneurons, or to rescue KCC2 expression in principal cells. KCC2*Flag indicates shRNA-proof, Flag-tagged KCC2 sequence. **C.** Representative confocal maximal projection images of hippocampal coronal sections immunostained for GFP and KCC2 from mice infected with CaMKII-shNT or CaMKII-shKCC2-expressing vectors, showing massive KCC2 knockdown in the later. Scale: 500 µm. **D.** Representative confocal maximal projection images of area CA1 of hippocampal coronal sections immunostained for GFP, GAD67 and KCC2 from mice infected with vectors expressing mDlx-shmirNT or mDlx-shmirKCC2. Arrowheads show infected cells are GAD67+ interneurons and that mDlx-shmirKCC2 efficiently suppressed KCC2 expression in these cells, as evidenced by lack of somatic KCC2 immunostaining (crossed out arrows). Scale: 20 µm. **E.** Representative immunoblot of KCC2, GFP and Flag in hippocampal protein extracts from mice infected with U6-shNT or U6-shKCC2 and CamKII-KCC2*Flag expressing vectors showing CamKII-KCC2*Flag efficiently restored KCC2 expression upon knockdown. **F.** Summary graph for contextual memory, showing % time freezing during the first 3 minutes of exploration of the foot-shock associated cage in a fear-conditioning paradigm, showing reduced freezing in mice infected with CaMKII-shmirKCC2 expressing vector as compared with mice infected with CaMKII-shmirNT vector (t-test, *p=0.019 for CaMKII-shmirKCC2). Mice infected with mDlx-shmirKCC2 vector or with U6-shKCC2 and CamKII-KCC2*Flag show no alteration of freezing behavior as compared to control mice. **G.** Summary graphs showing cued memory retention with no difference between groups.

These results show that KCC2 silencing only in the dorsal hippocampal principal neurons, but not interneurons, is sufficient to impair contextual fear memory. In order to further test the specificity of our knockdown approach, we then performed a rescue experiment in which KCC2 silencing was induced in dorsal hippocampus using an AAV1-pU6-shKCC2-CMV-GFP vector and then specifically rescued in principal cells using an AAV1-pCaMKII-KCC2*Flag vector, expressing a Flag-tagged and shRNA-proof recombinant KCC2 (Figure 2B). Co-infection was confirmed by western blot analysis with GFP and Flag antibodies (Figure 2E). Recombinant KCC2 overexpression only in principal cells fully restored contextual memory in mice with cell-type unspecific KCC2 silencing (Mann-Whitney rank sum test p=0.402; Figure 2F). Again, none of these manipulations induced significant change in anxiety (Figure EV1B-D) or locomotor activity (Figure EV1-F), as evaluated in open field and elevated-O-maze tests.

### KCC2 silencing enhances neuronal excitability and impairs hippocampal rhythmogenesis

Memory consolidation relies in part on hippocampal rhythmic activities (Colgin, 2016), such as theta- and gamma-band activities that occur during rapid eye movement (REM) sleep (Boyce *et al*, 2016; Buzsaki, 2002), as well as sharp-wave ripples (SWRs) associated with non-REM sleep and immobility (Buzsaki, 2015; Girardeau *et al*, 2009). Such activities involve temporally precise interactions between glutamatergic and GABAergic neurons (Adamantidis *et al*, 2019; Amilhon *et al*, 2015; Colgin, 2015; Gan *et al*, 2017; Valero *et al*, 2015). Since KCC2 knockdown is known to affect both glutamatergic (Gauvain et al 2011; Chevy et al 2015) and GABAergic (Goutierre *et al*., 2019; Pellegrino *et al*, 2011; Rivera *et al*., 1999) synaptic function as well as neuronal membrane excitability (Goutierre et al 2019), we hypothesized that these combined effects may impair hippocampal rhythmogenesis. To test this hypothesis, we bilaterally injected mice with AAV1-pU6-shNT or AAV1-pU6-shKCC2 vectors in the dorsal hippocampus and then implanted a silicon probe in unilaterally in one hippocampus (Figure 3A-C). The silicon probe was positioned to record LFP signal throughout the CA1 to DG axis.

**Figure 3.**
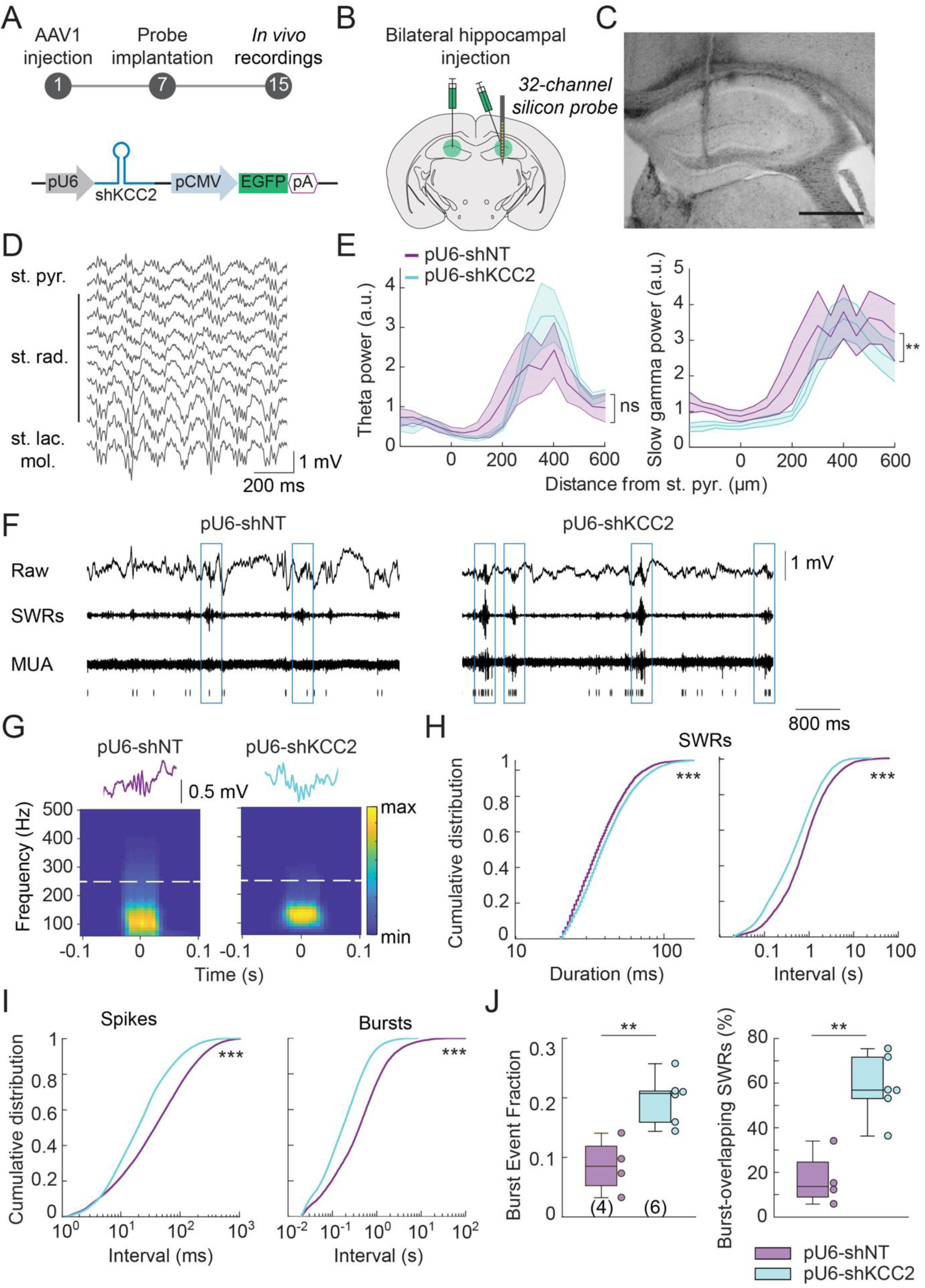
KCC2 downregulation leads to hippocampal network hyperexcitability. **A.** Timeline of the experiment (days) and viral vector used for KCC2 knockdown. **B.** Following the bilateral viral injection to knockdown KCC2 in dorsal hippocampus, a 32-channel linear silicon probe was implanted. **C.** Widefield macroscope image of dorsal hippocampus near implantation site showing the track of the implanted probe through CA1 and the dentate gyrus. Scale: 500 µm. **D.** Representative example of REM sleep recordings in mice infected with U6-shNT expressing vector. **E.** Left, theta power in the 5-10 Hz band measured during REM sleep using multi-tapers estimates. Power (in arbitrary units, a.u.) is plotted as a function of the electrode localization with respect to *st. pyramidale*. Right, theta power profile was not affected upon KCC2 knockdown (Kruskal-Wallis test, p=0.774). Slow gamma (25-55 Hz) power plotted as a function of the electrode localization with respect to *st. pyramidale*. Slow gamma power was reduced upon KCC2 knockdown (Kruskal-Wallis test, **p=0.003). **F.** Representative examples of LFP signal recorded in CA1 *st.pyramidale* mice infected with U6-shNT or U6-shKCC2 expressing vectors during non-REM sleep. The signal was filtered to detect either sharp wave ripples (SWRs, boxed in blue, 100-300 Hz) or multiunit activity (MUA, >500 Hz), as indicated in raster plot below. **G.** Top, representative recordings of individual ripples from mice infected with U6-shNT or U6-shKCC2 expressing vectors. Bottom, time-frequency plots showing the average frequency over time of all ripples in recordings from 4 control and 5 KCC2 knockdown mice. Note the lack of high-frequency oscillations or fast ripples (>250 Hz) in either condition. **H.** Cumulative distribution functions of ripple duration (left) and inter ripple interval (right). Upon KCC2 knockdown, both ripple duration and frequency significantly increased (Kolmogorov–Smirnov test, ***p<0.001). **I.** Cumulative distribution functions of inter-spike (left) and inter-burst (right) intervals. MUA frequency as well as burst frequency were increased in KCC2-knockdown mice compared to controls (Kolmogorov–Smirnov test, ***p<0.001). **J.** Summary boxplots showing the distribution of burst event fraction (number of bursts relative to the number of spike trains (left) and proportion of ripples associated with bursts of MUA activity (right). Both were significantly increased upon KCC2 knockdown (Mann-Whitney, **p=0.010).

Theta-band activity (5-10 Hz) was recorded throughout hippocampal layers during REM sleep, with maximal power within *st. lacunosum moleculare*, as previously reported (Buzsaki, 2002; Goutierre *et al*., 2019) (Figure 3D-E). No difference was observed in the power profile of theta-band activity between KCC2 knockdown and control mice (Kruskall-Wallis test p=0.774). Gamma-band activity is also detected during various behavioral states including REM sleep and can be divided in slow (25-55 Hz) and fast (60-90 Hz) gamma components, with distinct underlying mechanisms and functional impact on memory encoding and consolidation (Colgin, 2015, 2016). We therefore distinguished the two components and observed a significant and specific decrease in the power of slow (Kruskall-Wallis test p=0.003, Figure 3E) but not fast gamma oscillations (p=0.247) in KCC2 knockdown mice as compared to control.

We then tested whether KCC2 silencing might affect SWRs during slow-wave sleep (Figure 3F-H). Whereas SWR duration increased modestly (by about 10 %) in KCC2 knockdown as compared to control mice, their mean rate increased by 140% (Figure 3H; Kolmogorov-Smirnov test p<0.001 for both). However, their time-frequency profiles were similar, with no sign of high-frequency oscillation or fast-ripple component (250-500 Hz) that represent hallmarks of the epileptic hippocampus (Levesque *et al*, 2017; Menendez de la Prida & Trevelyan, 2011)(Figure 3H). This lack of epileptiform activity was consistent with the lack of spontaneous seizures in KCC2 knockdown animals observed during weeks of behavioral evaluation.

More strikingly, however, we observed a large and significant increase (by about 125 % on average) in multiunit activity (MUA) within *st. pyramidale* of CA1 in KCC2 knockdown mice as compared to controls (Figure 3F,I). This increase in frequency (Kolmogorov-Smirnov test p<0.001, Fig. 3I) was not homogenous but was associated with a remarkable (+268 %) increase in the occurrence of bursts of MUA (Kolmogorov-Smirnov test p<0.001; Figure 3F,I). This increased burst frequency was not just a mere reflection of the increased MUA, as the burst event fraction (number of bursts divided by the number of spike trains) was also significantly increased in KCC2 knockdown mice (Figures 3J; Mann-Whitney rank sum test p=0.010). Moreover, spike bursts were more often associated with ripples than in control mice (Figure 3N, Mann-Whitney rank sum test p=0.010).

Altogether, these results show that KCC2 downregulation in dorsal hippocampus affects hippocampal rhythmogenesis by i) reducing slow gamma rhythm, ii) increasing SWR rate and duration and iii) increasing MUA and bursting in CA1 as well as promoting spike bursts during SWRs. Such neuronal hyperexcitability may then generate unspecific *noise* and compromise memory consolidation (Iwasaki & Ikegaya, 2021), thereby contributing to impair hippocampus-dependent memory.

### The specific Task-3 channel opener terbinafine reduces neuronal excitability and occludes memory impairment upon KCC2 silencing

KCC2 interaction with leak potassium channel Task-3 controls membrane trafficking and expression of Task-3, thereby influencing membrane resistance and excitability (Goutierre *et al*., 2019). Thus, KCC2 silencing was shown to increase hippocampal principal neuron excitability *in vitro*, independently of changes in GABAergic signaling. This effect promoted anomalous recruitment of dentate granule cells during dentate spikes which was prevented by chemogenetic silencing of granule cells *in vivo* (Goutierre *et al*., 2019). We hypothesized a similar effect may also contribute to increase neuronal firing and contamination of SWRs with bursts of action potentials upon KCC2 knockdown. We therefore tested whether increasing Task-3 function, using the selective activator terbinafine (Tian *et al*, 2019; Wright *et al*, 2017), might prevent neuronal hyperexcitability and thereby rescue or occlude memory upon KCC2 silencing. In KCC2-knockdown mice, we implanted an intracerebroventricular (icv) canula and a silicon probe to record neuronal activity in the right dorsal hippocampus while injecting terbinafine in the left lateral ventricle (Figure 4A). Infusion of terbinafine (20 µl, 500 mM solution) but not saline reduced multiunit activity in the CA1 area by about 40% for up to two hours (Figure 4C-E, t-test p=0.013).

**Figure 4.**
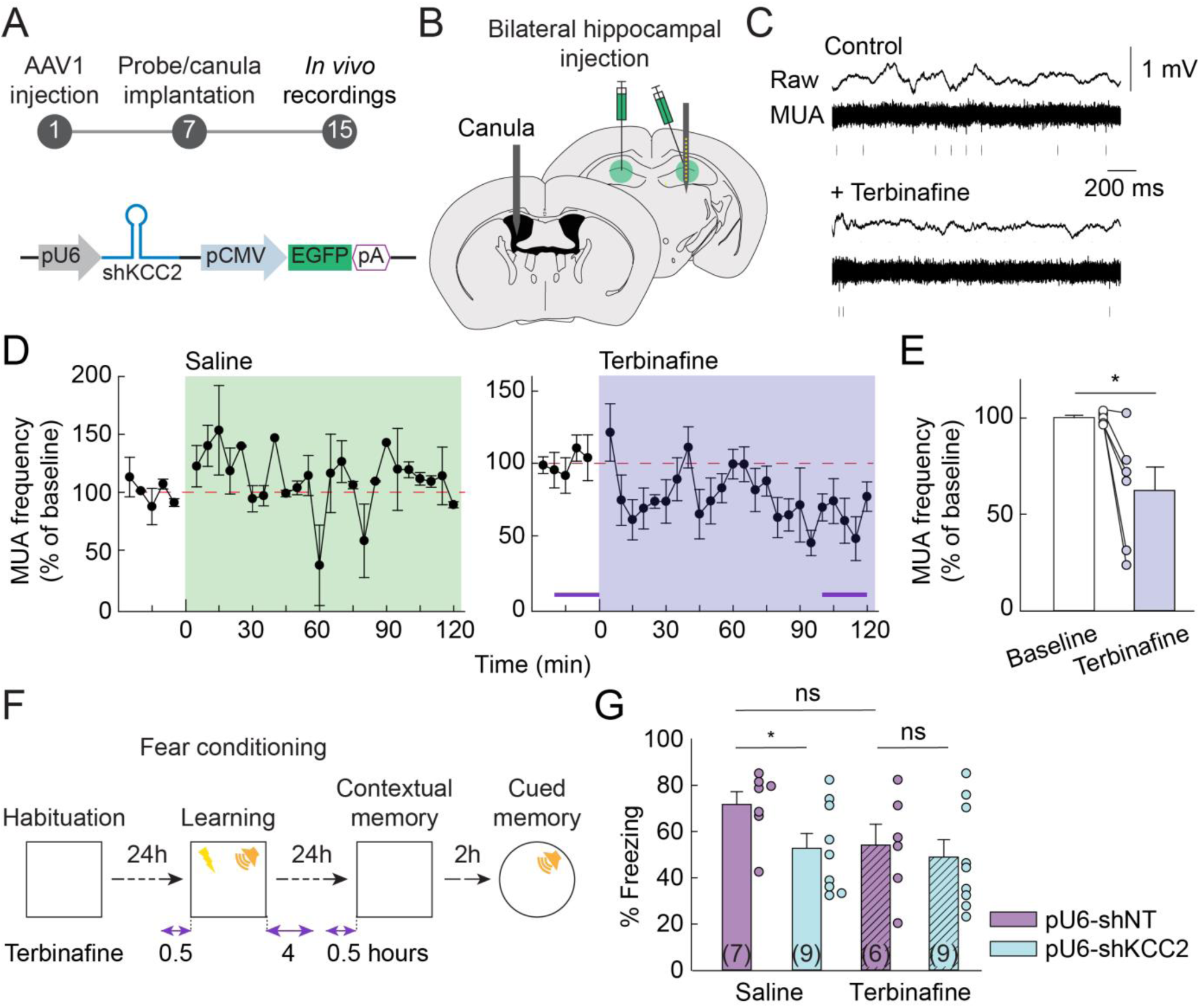
Task-3 channel activator terbinafine occludes memory deficits upon KCC2 knockdown. **A.** Experimental timeline (days) and viral vector used in these experiments. **B.** A week after bilateral hippocampal injection of AAVs to knockdown KCC2, a silicon probe and a cannula were implanted in the right hippocampus and left lateral ventricle, respectively. **C.** Representative examples of LFP signals recorded in CA1 *st.pyramidale* of mice infected with U6-shKCC2 expressing vectors, before and after icv injection of terbinafine (20 µL of 500 mM solution). The signal was filtered to detect multiunit activity (MUA, >500 Hz). **D.** Summary plots showing MUA frequency over time, normalized to control (prior to icv injection) in animals infected with U6-shKCC2 vector, with saline (left, n=4 from 2 mice) or terbinafine (right, n=3 from 2 mice). **E.** Summary graph showing MUA frequency before and 110 min following icv terbinafine injection. (t-test, *p=0.013). **F.** Timeline of the fear conditioning paradigm with icv injection of saline or terbinafine. **G.** Summary plot showing % time spent freezing during 3 minutes of exploration of the foot-shock associated cage. Saline-injected mice infected with U6-shKCC2 vector display reduced freezing compared to control mice infected with U6-shNT expressing vector (t-test, *p=0.023). However, this effect was occluded by icv infusion of terbinafine (t-test, p=0.346). Terbinafine induced a noticeable yet non-significant reduction of freezing in the control group (U6-shNT, t-test, p=0.06).

We then tested the effect of intraventricular terbinafine infusion on contextual memory in mice following KCC2 silencing. Mice were infected with either the pU6-shKCC2 or pU6-shNT viral vector and a canula was implanted to allow infusion of terbinafine or saline as a control. As in previous experiments, KCC2 knockdown mice receiving saline infusion showed reduced freezing as compared to control mice in a contextual fear memory paradigm (t-test p=0.023; Figure 4F-G). However, in mice receiving intraventricular terbinafine infusion, no difference in freezing was observed between KCC2 knockdown mice and controls (t-test p=0.104). Noticeably, control mice receiving terbinafine infusion showed reduced freezing levels compared to saline-treated mice (t-test p=0.056), suggesting intraventricular terbinafine treatment may also partially impair contextual memory. Interestingly, infusion of terbinafine did not further affect contextual memory in KCC2 knockdown mice. Together, our results show that KCC2 silencing induces neuronal hyperexcitability in dorsal hippocampus and that reducing excitability with a Task-3 channel activator is sufficient to occlude impairment of contextual memory.

## Discussion

We show that chronic KCC2 downregulation in the dorsal hippocampus induces spatial and contextual memory deficits that can be mimicked by KCC2 silencing in principal neurons but not GABAergic interneurons. This memory impairment is associated with neuronal hyperexcitability and bursting as well as altered hippocampal rhythmogenesis, but no detectable spontaneous epileptiform activity. Finally, our results show that intraventricular administration of terbinafine, a specific Task-3 channel activator, reduces neuronal hyperexcitability and occludes contextual memory impairment upon KCC2 silencing in dorsal hippocampus.

Reduced neuronal KCC2 expression has been reported in animal models of epilepsy (Kourdougli *et al*, 2017; Pathak *et al*., 2007) as well as postoperative tissue from intractable epilepsy patients (Blauwblomme *et al*, 2019; Huberfeld *et al*., 2007; Munakata *et al*, 2007; Pallud *et al*., 2014; Palma *et al*, 2006). Computational modeling predicted that complete KCC2 suppression in only 30% of principal neurons may be sufficient to promote seizures, at least under conditions of neuronal hyperexcitability (Buchin *et al*., 2016). Here, this hypothesis was tested experimentally and show that chronic KCC2 silencing throughout mouse dorsal hippocampus affects neuronal and network activity but fails to trigger detectable epileptiform activity or seizures. Similarly, no sign of an epileptic network was detected upon KCC2 knockdown in rat dentate gyrus (Goutierre *et al*., 2019). These results contrast with those obtained upon complete genetic ablation of the KCC2b isoform (Woo *et al*, 2002) or conditional deletion of KCC2 carboxy-terminal domain in mouse dorsal hippocampus (Kelley *et al*, 2018), both leading to spontaneous seizures. This suggests that the complete loss of KCC2 expression or expression of a truncated, likely dysfunctional transporter may induce a more severe phenotype than that induced by KCC2 downregulation as observed in the pathology. Further supporting this conclusion, hypomorphic KCC2-deficient mice retaining 15-20% of normal KCC2 expression also failed to display spontaneous seizures (Tornberg *et al*, 2005).

Our data therefore demonstrate that memory deficits are induced by KCC2 downregulation independent of epileptiform activity. Memory involves a sequence of events (encoding, consolidation and retrieval, (Bliss & Collingridge, 1993; Colgin, 2016)) which may all, in principle, be affected by KCC2 downregulation. Expression of long-term potentiation, a key mechanism for engram formation and selection (Kim & Cho, 2017; Nabavi *et al*, 2014), is hindered upon KCC2 knockdown in hippocampal neurons due to altered activity-dependent AMPA receptor trafficking in dendritic spines (Chevy *et al*., 2015). Impaired hippocampal LTP may thus contribute to the spatial and contextual memory deficits upon KCC2 downregulation in dorsal hippocampus. In addition, hippocampal rhythmogenesis is involved in both memory encoding and consolidation (Colgin, 2016). Given the importance of GABA signaling in both hippocampal theta- and gamma-band oscillations (Csicsvari *et al*, 2003; Wulff *et al*, 2009), which are critical to memory encoding and retention (Colgin, 2016; Hasselmo, 2005), it is remarkable that KCC2 downregulation had only little effect on these oscillations, with only a moderate reduction of slow gamma band activity power. This may reflect the relative preservation of the driving force of GABAA receptor-mediated currents upon KCC2 knockdown, due to the parallel depolarization of their reversal potential and of resting membrane potential (Goutierre *et al*., 2019). One of the most striking alteration of hippocampal activity induced upon KCC2 knockdown was increased neuronal firing and bursting. This effect was particularly apparent during non-REM sleep, where bursts of multiunit activity often superimposed with SWRs, which play a crucial role in memory consolidation (Buzsaki, 2015; Girardeau *et al*., 2009), and was associated with an increase in their duration. During SWRs, co-firing patterns of neurons activated during previous wakefulness are replayed (Pfeiffer, 2020), likely contributing to memory consolidation. Neuronal hyperexcitability and bursting resulting from KCC2 knockdown may then act to scramble the information content of replay firing sequences, thereby compromising the specificity of memory consolidation. Nonspecific firing of just few hippocampal CA1 neurons was indeed shown to compromise reactivation of memory engram cells (Iwasaki & Ikegaya, 2021). Increased hippocampal activity correlating with cognitive decline has been reported in aging rodents and can be rescued by using low doses of antiepileptic drugs acting to decrease neuronal excitability (Bakker et al., 2012; Koh et al., 2010; Wilson et al., 2005). Similarly, here, normalizing neuronal activity with a Task3-specific channel opener occluded the contextual memory deficits induced by KCC2 suppression, suggesting neuronal hyperexcitability may play a critical role in these deficits.

Together, our results demonstrate that KCC2 downregulation, as observed in numerous neurological and psychiatric disorders, leads to neuronal hyperexcitability and memory impairment and that Task-3 channels openers, like KCC2 enhancers (Gagnon *et al*, 2013; Tang *et al*., 2019), may be of therapeutic interest for cognitive deficits associated with these disorders.

### Materials and Methods

### • RNA interference and viral vectors

In order to suppress KCC2 expression in mice, we used RNA interference with a previously validated short hairpin RNA (shRNA) sequence (shKCC2: AGCGTGTGACAATGAGGAGAACTTCCTGTCATTCTCCTCATTGTCACACGCT, (Bortone & Polleux, 2009)). A sequence devoid of target in mouse genome (shNT: GGAATCTCATTCGATGCATACCTTCCTGTCAGTATGCATCGAATGAGATTCC, (Gauvain *et al*., 2011)) was used as control. These sequences were inserted in pAAV vectors under U6, CaMKII or mDlx promoters. Since the two later are polymerase II promoters, shRNA sequences were then embedded into micro-RNA (mir30) backbone (Fellmann *et al*., 2013). Finally, once plasmids were cloned (pAAV-U6-shKCC2(shNT)-CMV-GFP-SV40, pAAV-CamKII-shmirKCC2(shmirNT)-SV40-CMV-GFP-SV40, pAAV-mDlx-GFP-shmirKCC2(shmirNT)-WPRE-SV40), they were used to produce purified AAV particles for co-expression with GFP (AAV2.1, titer 10^13^ TU/ml, Atlantic Gene Therapy, Nantes). For KCC2 expression rescue, a Flag-tagged recombinant KCC2 sequence (Chamma *et al*, 2013) was modified to make it shRNA-proof. Mutated shRNA target sequence was synthetized (Genscript) and then subcloned in place of the original target sequence using classical digestion, ligation and transformation. Sequence was mutated in order to prevent shRNA complementarity without changing amino-acid sequence (AACGAGGTCATCGTGAATAAATCC modified to AATGAAGTGATTGTCAACAAGTCC). The sequence was cloned into a pAAV-CamKII-KCC2flag-WPRE-SV40 vector to replace the KCC2flag sequence and used to produce purified AAV particles as above.

### • Animals

C57Bl/6JRj mice were housed in standard laboratory cages on a 12-hours light/dark cycle, in a temperature-controlled room (21°C) with free access to food and water. Mice were purchased from Janvier Labs (Le Genest-Saint-Isle, France) and were delivered to our animal facility at least a week before surgery or behavioral testing. All procedures conformed to the International Guidelines on the ethical use of animals, the French Agriculture and Forestry Ministry guidelines for handling animals (decree 87849, license A 75-05-22) and were approved by the Charles Darwin ethical committee (agreement 2016042916319074v5).

### • Stereotaxic surgery

Mice were anesthetized with ketamine/xylazine (100/15 mg/kg) and placed on a heating pad at 36-37°C for the entire surgery. 10 minutes before opening the skin, lidocaine (2%) was applied locally on the skin. The virus was bilaterally injected in dorsal hippocampus (500 nl in both the dentate gyrus and CA1 for each hemisphere) at the following stereotaxic coordinates from Bregma: -1.8 mm anteroposterior (AP), +/- 1.2 mm mediolateral (ML) and - 2.1/-2.0/-1.9/-1.3/-1.25 mm dorsoventral (DV). After surgery, body temperature was maintained using a heating pad under the cage until the animal recovered from anesthesia. The analgesic carprofen (0.5 mg/mouse/day) was then added to the drinking water for the next 24 hours. Behavioral experiments or electrophysiological recordings were then conducted 10 to 14 days after surgery.

### • Silicon probe implantation

One week after viral injection, mice were implanted with a 16 or 32-channel linear silicon probe (Neuronexus A1x16-5mm-100-413-CM16LP or A1x32-6mm-50-177-CM32). Mice were anesthetized with isoflurane (4 % for induction, 1-2 % for maintenance) in a stereotaxic frame for the entire surgery and their body temperature was maintained with a heating pad. In order to reduce pain, mice were injected with buprenorphine before surgery (0.1 mg/kg). First, the skull was cleaned and the craniotomy at the probe location was re-opened. Two screws were implanted on the frontal bone (1 per hemisphere), 1 on the parietal bone (contralateral to the probe) and 1 reference screw above the cerebellum. Then, the probe was descended into the dorsal right hippocampus at the following coordinates from Bregma: -1.8 mm (AP), -1.2 mm (ML), -2.4 mm (DV, for the 32-channels silicon probes) or - 2.0 mm (DV, for the 16-channels silicon probes). Once the probe was positioned, the craniotomy was covered with Vaseline to protect the probe.

The head-stage was then built using a thin layer of SuperBond dental cement applied onto the skull, followed with Unifast TRAD dental cement on the probe and the screws. Pieces of copper mesh were then arranged around the probe to create a Faraday cage and cemented onto the skull after soldering the ground and reference of the probe with the reference electrode above the cerebellum to the copper mesh.

After surgery, body temperature was maintained with a heating pad under the cage until the animal recovered from anesthesia. The mice were then housed separately to avoid damage to the probe. For 3 days after surgery, mice were fed with high-calorie liquid chocolate and received two daily injections of buprenorphine (0.1 mg/kg i.p.). Behavioral experiments and recordings were performed one week after surgery, once the animals had fully recovered.

### • Canula implantation and terbinafine treatment

A home-made canula (Kokare *et al*, 2011) was implanted into the left ventricle at the following coordinates from Bregma: -0.4 mm (AP), -1.0 mm (ML), -3.0 mm (DV), either on the day of silicon probe implantation or the day of viral injection. Mice were handled daily for a week and received 10 µL of salin**e** injection to habituate them to intracerebroventricular (icv) injection. On the fear conditioning day, mice received 20 µL of 500 mM terbinafine solution in saline or 20 µL of saline, 30 minutes before experiencing the tone-shock association. They received a second injection 4 hours later. On the next day, mice received a third injection 30 minutes before testing contextual memory retrieval.

### • *In vivo* recordings

Animals implanted with silicon probes were connected to a recording controller (Intan Technologies, Los Angeles, USA). 30 minutes of sleep were recorded and analyzed after 10 minutes exploration of an arena. Data were acquired at 20 kHz using Intan Recording Controller software (version 2.05). All analyses were performed offline using Matlab built-in functions, Chronux (Bokil *et al*, 2010), the FMAToolbox (http://fmatoolbox.sourceforge.net/), as well as custom-written scripts.

Power spectra and spectrograms were computed using multi-tapers estimates on the raw LFP signal. Gamma-band was defined as 25-90 Hz. Slow and fast gamma ranges (respectively 25-55 and 60-90 Hz) were determined according to (Colgin *et al*, 2009). Theta power was determined in the 5-10 Hz band. Spike detection was performed using high-pass (>500 Hz) filtering and semi-automatic thresholding. Bursts of spikes were defined as a minimum of 3 spikes with inter-spike interval less than 20 ms. This value was determined from the distribution of all inter-spikes intervals (ISI) where 40-50 % recordings show ISI <20 ms in our data set. Ripple detection was performed by band-pass filtering (100-600 Hz), squaring and normalizing, followed by thresholding of LFP recorded in CA1 pyramidal layer. Thresholds were different between each animal and were determined manually, in order to detect only SWRs.

### • Behavioral tests

#### Open field

Mice were gently placed facing the wall of a large white arena (50 x 50 x 40 cm, 90-100 lux) for a 10 minutes recording period. Their activity was recorded and analyzed using Ethovision software (Noldus). Mice were considered in the center of the arena (1/9 of the arena) when their body center was in the central zone. The arena was cleaned with 10% ethanol.

#### Elevated-O-maze

The apparatus consisted in a white circular maze (35 cm of diameter, lane is 6 cm wide), elevated 50 cm above the floor, and divided in 4 quadrants: two opposite “closed” quadrants surrounded by 12 cm high opaque walls (the safe environment) and two “open” quadrants. The luminosity was set at 8-10 lux in the open arms and the elevated-O-maze was cleaned with 10% ethanol. Mice were placed randomly in one of the closed arms, facing an open arm. Their activity was recorded for 10 minutes and analyzed using the Ethovision software. Mice were considered in the open arms when the four paws were placed in the arm.

#### Place recognition

The arena and the objects were cleaned with 10% ethanol and the luminosity in the center of the arena was set to 10 lux in order to promote exploration. A cue card was placed on one wall to allow to orient the mice in the arena. On the three first days, mice were habituated to the arena, with two habituations per day, at least 3 hours apart. On the first habituation (H1), all mice from a single cage were placed in the arena for 30 minutes, allowing them to explore together the environment to help reduce stress. For the next four habituations (H2 to H5), mice were placed alone in the arena for 10 minutes. On the last habituation (H6), two identical objects were introduced to help reduce stress associated with novelty.

During the first exposure, 24 to 72 hours after H6, two identical objects were presented to the mouse for 10 minutes. Then, 10 minutes (short-term memory) or 24 hours later (long-term memory), mice were placed back in the arena with one object moved and the other in the same position. Mouse behavior was video-recorded and analyzed manually during the exposure session, in order to ensure that mice were exploring enough and did not have an initial preference for one of the objects. The data recorded during the memory test were manually scored. A discrimination index (DI) was used as a measure of discrimination between novel and old location and obtained by dividing the difference in time spent in the old versus new locations by the total exploration time. Exploration was considered only when the animal was both in contact and facing the object.

#### Fear conditioning

On day 1, mice were exposed to the experimental chamber and let free to explore. Each mouse was placed four minutes in box 1 (27x27 cm, black environment, white light), cleaned with 70% ethanol. On the second day, mice were allowed to explore box 1 for 4 minutes before a tone was delivered for 30 seconds (4 Hz, 85 dB) that co-terminated with a two-seconds foot shock (0.25 mA) delivered through the floor grid. After a 30-second interval, the tone-foot shock procedure was repeated. On the third day, mice were placed back in box 1 for 4 minutes to evaluate contextual memory (i.e., without receiving any foot-shock). Two hours later, they were placed in a box 2 to test cued memory. Box 2 was white and round, with no grid on the floor, red light, and was cleaned with 1% acetic acid. After 4 minutes of exploration, the tone (4 Hz, 85 dB) was played for 30 seconds, followed by 30 seconds of silence. This sequence was repeated 6 times.

The freezing behavior was recorded and analyzed either with Packwin v2.0.05 software (Panlab) or manually.

### • Western blot analysis

Protein concentration in all samples was determined using the Pierce assay (Pierce™ BCA Protein Assay Kit, Thermo Fisher). Equal amounts of proteins were then separated on a 4%–12% SDS polyacrylamide gradient gel (Invitrogen) and transferred onto a nitrocellulose membrane (GEHealthcare) at 100 V for 1 hour in order to ensure efficient transfer of high molecular weight proteins using a liquid transfer system (Bio-Rad). The membrane was blocked for one hour at room temperature in 5% milk (diluted in 1X TBS – 0,1% Tween 20) then washed three times for 5 min in 1X TBS – 0,1% Tween 20. These washing conditions were used after incubation with primary and incubation with secondary steps. The membrane was incubated overnight at 4°C with the primary antibody (diluted in 5% milk, 1X TBS – 0,1% Tween 20). Then the secondary antibody (diluted in 5% milk, 1X TBS – 0,1% Tween 20) was added for 1 hour at room temperature. The membrane was then probed using the Odyssey blot imaging system (LI-COR).

The primary antibodies used were: rabbit anti-KCC2 (1:1000, Millipore), mouse anti-Tuj1 (1:10000, R&D systems). The secondary antibodies used were: goat anti-rabbit 800 (1:5000, Tebu-Bio or 1:15000, LiCor), goat anti-mouse 700 (1:5000, Tebu-Bio or 1:15000, LiCor).

### • Histology

Mice were deeply anesthetized using sodium pentobarbital (200 mg/kg i.p.) and then transcardially perfused with ice-cold PBS solution, followed by ice-cold 4% paraformaldehyde (PFA) in PBS pH 7.4. Extracted brains were fixed in 4% PFA overnight and then equilibrated in 30% sucrose in phosphate buffer saline (PBS). A sliding cryotome was used to section the brains into 40 μm-thick coronal sections. Immunohistochemistry experiment was performed when necessary. Only mice with a minimal infection spread of 500 µm in both dorsal hippocampi (CA1, CA3, DG) were considered for further behavioral analysis.

### • Immunohistochemistry

Slices were incubated for 1 hour at room temperature in a blocking solution containing 10% goat serum, 0.1% Triton X-100 in PBS. Incubation with primary antibody (rabbit anti-KCC2 (1:500, Millipore), chicken anti-GFP (1:1000, Millipore), mouse anti-Flag (1:1000, Sigma-Aldrich), mouse anti-GAD67 (1:1000, Abcam)) was then performed in blocking buffer overnight at 4°C. After 3 x 10 min washes in PBS, slices were incubated with secondary antibodies with a dilution of 1:1000 in blocking buffer for 1 hour (goat anti-rabbit-cy3, donkey anti-mouse-cy5, donkey anti-chicken-488 (Jackson Labs)). After another 3 x 10 min washes, slices were mounted on a coverslip in Mowiol/Dabco (25 mg/mL) solution. Immunofluorescence images were acquired using an upright confocal microscope (Leica TCS SP5), using a 40X 1.30-N.A. objective and Ar/Kr laser set at at 491 and 561 nm for excitation of Cy3 and FITC, respectively, for images of the entire hippocampus. Stacks of 25-30 µm optical sections were acquired at 512x512 pixel resolution with a z-step of 1 µm.

### • Statistics

All statistical tests were performed using SigmaPlot software (Systat Software Inc.). When necessary, proportions were arcsin-transformed prior to performing appropriate statistical test. Comparison of means was performed using Student’s t-test for normally distributed variables (as tested with Shapiro-Wilk test) of equal variances (Brown-Forsythe test). Otherwise, comparison of mean was performed using the non-parametric Mann-Whitney rank sum test. Two-way ANOVA was used for comparison of distributions when data were normally distributed with equal variances. Kolmogorov-Smirnov test was used to compare distributions. Statistical significance was set to p≤0.05.

## Acknowledgements

We thank Liset M de la Prida and Gabrielle Girardeau for helpful discussions and critical reading of the manuscript. We acknowledge the Imaging Platform of Institut du Fer à Moulin, where confocal imaging was performed and Atlantic Gene Therapy (UMR-1089, Univ. of Nantes) for AAV production. This work was supported in part by Fondation pour la Recherche Médicale (DEQ20140329539 to JCP), ERANET-Neuron (funded by Agence Nationale de la Recherche to JCP) and the Fondation Française pour la Recherche sur l’Epilepsie - Fédération pour la Recherche sur le Cerveau (to JCP). C.S. and M.G. were recipients of fellowships from Sorbonne University and C.S. was partly supported by the Bio- Psy Laboratory of Excellence. The Poncer lab was affiliated with the Paris School of Neuroscience (ENP) and the Bio-Psy Laboratory of Excellence.

## Author contributions

C.S., S.D. and J.C.P. designed the research; C.S. and M.S. performed the experiments; C.S. and M.S. analyzed the data; M.G. and M.S. wrote the Matlab codes for data analysis; I.M. designed vectors for virus production; S.D. and J.C.P. supervised the research; C.S. and J.C.P. prepared the figures and wrote the manuscript with inputs from M.S. and S.D.

